# Exceptional diversification of floral form in a specialized orchid pollination system

**DOI:** 10.1101/2025.07.25.666443

**Authors:** Jasen W. Liu, Diego Bogarín, Oscar A. Pérez-Escobar, Franco Pupulin, Adam P. Karremans, Zuleika Serracín, Yongxuan Xie, Eugenio Restrepo, Santiago R. Ramírez

## Abstract

Traits that facilitate specialized interactions, such as those in flowers that promote pollination, are often invoked as targets of stabilizing selection across macroevolutionary timescales. However, the diversity of pollination mechanisms across flowering plants begs further investigation into the generality of this pattern. We fit a model of multivariate character evolution on a dataset of 140 orchid species sampled across 65 genera from the diverse neotropical Cymbidieae clade to characterize the role of pollination mode on the pace of flower shape evolution. We find that, contrary to the expectation of pollinator-mediated stabilizing selection causing stasis, orchids pollinated by specialized scent-collecting male euglossine bees (“perfume flowers” sensu (1, 2)) exhibit elevated rates of floral evolution compared to plants utilizing other rewarding or deceptive mechanisms. This pattern is recapitulated across at least 5 independent origins of this pollination system amidst a complex backdrop of background rate evolution. The rapid rates of change we observed in perfume flowers may be facilitated by weak evolutionary coupling between functional regions in their flowers, allowing for independent trajectories of evolution. Our results provide novel insights into the capacity for pollinators to generate selective pressures on flowers at macroevolutionary scales, providing an engine for trait diversification in some of the world’s most floristically rich regions.

## Introduction

Convergence and stasis are widely regarded as the key influences of specialized pollinators on floral evolution, driving stabilizing selection across macroevolutionary timescales (3, 4). A considerable body of literature has documented patterns of covarying suites of floral traits in relation with certain pollinator functional groups – “pollination syndromes”(5, 6). Strong patterns of convergence of floral form are seen coincident with transitions to certain pollinator groups that allow for efficient pollination, such as the repeated evolution of long nectar spurs and phenylpropanoid-dominated scents in hawkmoth-pollinated “moonflowers” (7, 8). Furthermore, lineages exhibiting certain pollination modes have been found to exhibit patterns of morphological stasis due to persistent selective forces on floral form that best mediates pollinator interactions (3, 4, 9). However, thus far the ability of pollinators to exert selection on macroevolutionary timescales to affect the tempo of floral morphological evolution of the plants they pollinate is unclear (6).

Orchids of the neotropical clade Cymbidieae offer a powerful system to test macroevolutionary hypotheses about trait diversification mediated by pollinators. The roughly 3400 species of this clade originated from an Australasian dispersal event into tropical America around 34 million years ago (10). In their subsequent radiation they have evolved a myriad of pollination mechanisms and reward strategies. Critically, there have been several independent origins of a highly specialized pollination system – androeuglossophily (11). These flowers are exclusively pollinated by male euglossine bees that are attracted by scent and collect these chemical rewards for use in species-specific courtship displays (1, 2, 12, 13). As such, scent operates as both a signal and reward in this pollination mutualism, differentiating its function from other attraction mechanisms where scent either signals the presence of a separate reward or is used to exploit sensory biases of pollinators expecting rewards in deceptive flowers. The ease of androeuglossophily in generating reproductive isolation via simple changes in scent production is thought to increase speciation rates in orchids (14). While prolific chemical convergence occurs in the scent of these plants across different scales of taxonomic organization (15, 16), the influence of this specialized pollination mechanism on morphological evolution is not well characterized.

As scent acts simultaneously as a signal and a reward in perfume flowers, selection on morphological elements of visual cues may be relaxed. Male euglossine bees readily collect scents from diverse sources outside of orchid flowers such as aromatic leaves, feces, and even filter paper soaked with pure compounds, indicating a lack of context-dependent visual cues to elicit such behavior (17–19). Consequently, there may be reduced constraints on the evolution of morphologies mediating precise physical fit of pollinators with the floral reproductive apparatus. These diverse mechanisms allow for different species of perfume flowers to place their pollen structures on different parts of bee bodies, partitioning the over 40 species of bees found within many communities (20, 21). Perfume flowers utilize some of the most complex pollination mechanisms in the plant kingdom, often involving the directed movement of pollinators through their flowers to achieve precise fit with reproductive organs – for example, escaping a “bucket” of liquid through a narrow tunnel containing the reproductive organs in *Coryanthes*, or the forceful deposition of pollinaria across the genus *Catasetum* (22). These specialized functional morphologies have captivated evolutionary biologists since the dawning of the field (23–25). Characterizing the evolutionary dynamics of variation in these traits in the context of recent phylogenetic reconstructions of orchids (10, 26) will allow for greater understanding of how pollinators shape floral evolution across macroevolutionary timescales.

Flowers are modular structures, consisting of distinct whorls of organs that mediate interactions with pollinator bodies to allow for efficient pollen transport and receipt. Phenotypes consisting of increased modularity (i.e. reduced covariation between structures) are often thought to facilitate diversification due to weaker constraints between the components allowing for independent evolutionary trajectories (27). Orchid flowers share a conserved bauplan with other lilioid monocots consisting of 3 petals and 3 sepals (Fig. S1). However, they additionally exhibit several lineage-specific novelties, most notably the modification of one petal into a “labellum” structure and the fusion of reproductive organs into a “column” structure. These modifications break the radial symmetry of the floral plan, creating a polarity amenable to the complex pollination mechanisms characteristic of this family. Labella have diversified greatly across the orchid radiation, usually acting to direct pollinators into a specific position to contact the column and facilitate pollination (28, 29). Understanding the degree to which the labella and column (henceforth referred to as the “reproductive structure”) and the unmodified petals and sepals (henceforth referred to as the “perianth”) are evolutionarily integrated will provide insight into understanding patterns of morphological evolution across the orchid floral phenotype.

Here we seek to address these gaps in knowledge by 1) characterizing the morphospace of neotropical Cymbidieae orchid flowers, 2) analyzing these morphological data within a phylogenetic context to understand the influence of pollination on the dynamics of orchid evolution at a macroevolutionary scale, and 3) characterizing patterns of correlation within and between different floral structures to understand how they are related to pollination and rates of evolution. Our results reveal that perfume flowers exhibit exceptionally high rates of morphological evolution and provide new insights into how pollinators shape the landscape of floral evolution across macroevolutionary timescales.

## Results

We characterized the morphology of the specialized column and labellum structures (“reproductive structure”) and unmodified petals and sepals (“perianth”) within the flowers of 178 species of orchids from the neotropical Cymbidieae, representing around 50% of genera and 100% of subtribes contained in this lineage. We found a broad diversity of floral morphologies in both sets of traits. The primary axis of morphological variation in the reproductive structure contrasts pouched labella, as found in the highly modified flowers of the “bucket orchid” *Coryanthes*, to highly reflexed labella, as exhibited by several species of *Cyrtochilum* (Figure 1A). The second major axis of variation describes a continuum from constricted tubes, formed by parallel positioning between the column and labellum, to structures where the labellum is positioned roughly 180 degrees from the column. The former phenotype is common among many species of the Maxillariinae (i.e. “gullet flowers” described in pollination biology literature of orchids (22)), while the latter phenotype is common among many species of the Oncidiinae, where the expanded flat labella operate as landing pads for bees seeking oil rewards (Fig. S2A).

**Figure 1:**
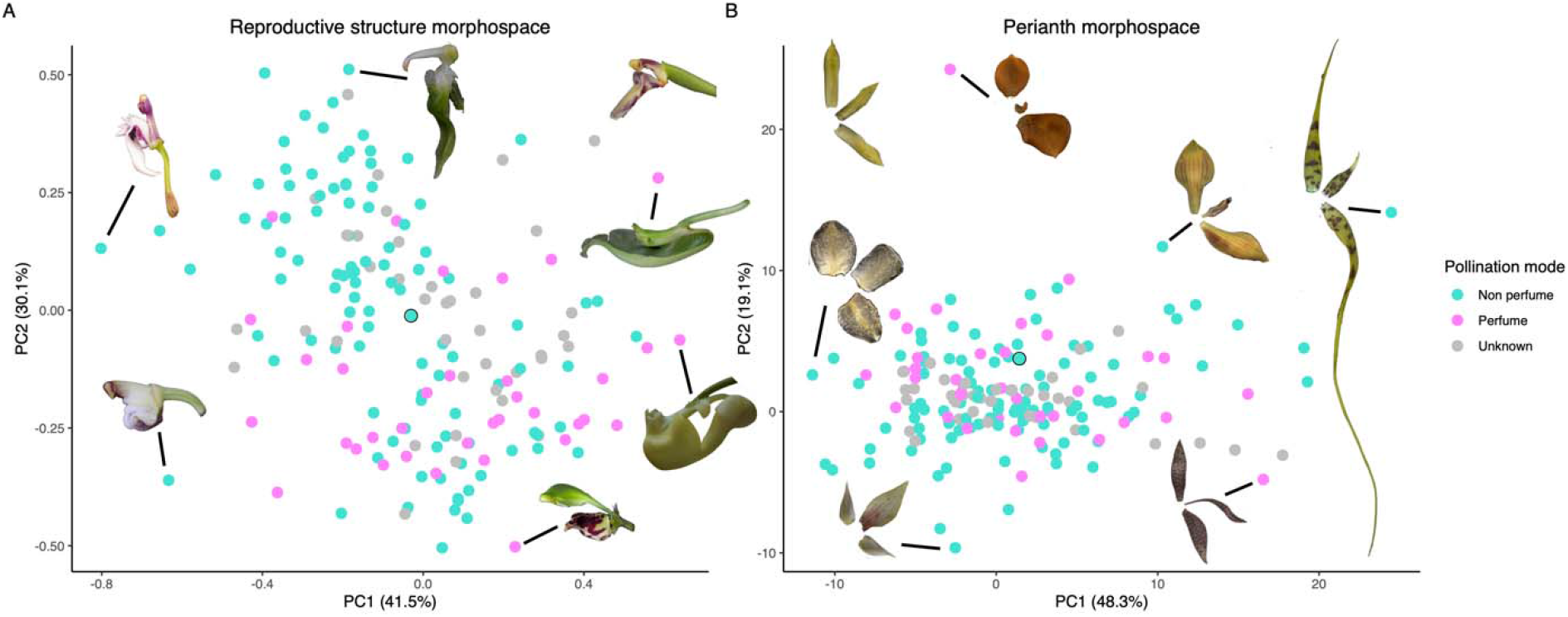
Morphospaces of the neotropical Cymbidieae flower comparing axes of variation in A. the reproductive structure and B. the perianth. Points represent species means and are colored by pollination mode while values on the x and y axis correspond to proportions of total morphological variance explained by the first and second principal components, respectively. Images on the top right of A and top left of B in boxes correspond to *Camaridium bracteatum*, a species close to the mean phenotype in both sets of traits (circled points in either panel). Other represented species in A are (from center above, going counterclockwise): *Ornithocephalus inflexus, Cyrtochilum ramosissimum, Cyrtochilum myanthum, Kegeliella kupperi, Coryanthes hunteriana*, and *Clowesia dodsoniana*. Other represented species in B are (from center above, going counterclockwise): *Gongora armeniaca, Trichocentrum morenoi, Telipogon biolleyi, Polycycnis barbata, Brassia caudata*, and *Trigonidium egertonianum*. Photographs copyright the authors except for the following, modified with permission: *Clowesia dodsoniana* from (55), *Cyrtochilum ramosissimum* from (56) copyright Carlos Jerez, *Cyrtochilum myanthum* from (57), and *Trichocentrum morenoi* copyright Gustavo Montealegre.

**Figure 2:**
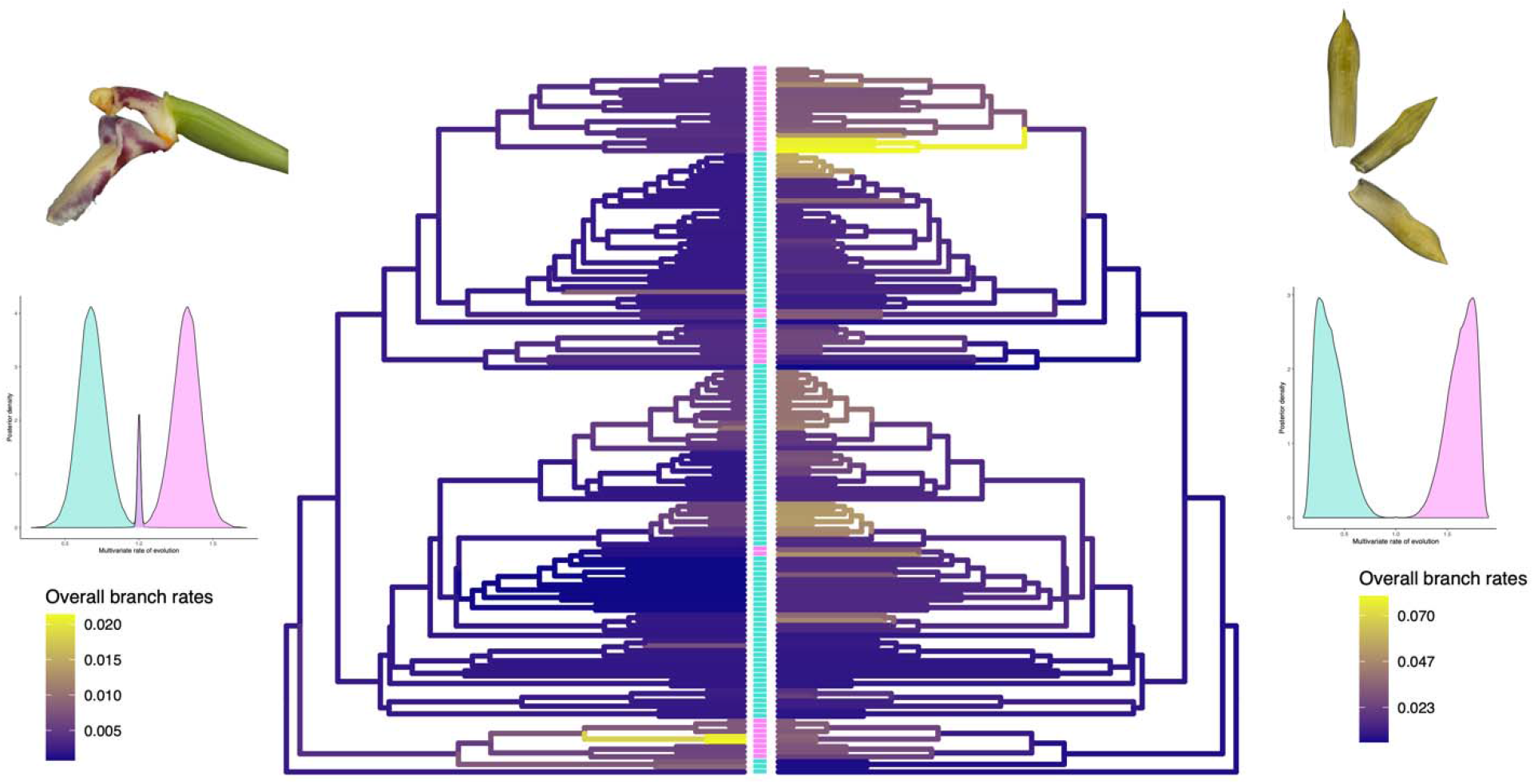
Rates of floral morphological evolution for the reproductive structure (left) and perianth (right) across the Cymbidieae radiation. Branch colors on the phylogenies indicate overall rates of evolution, with dark purple indicating lower rates and yellow indicating higher rates. Bars between the colored phylogenies represent character states at the tips, with pink corresponding to perfume flowers and turquoise corresponding to non perfume flowers. Plots below representative images of the floral structures represent posterior estimates of state-dependent evolutionary rates, with pink representing perfume flowers and turquoise representing non perfume flowers.

**Figure 3:**
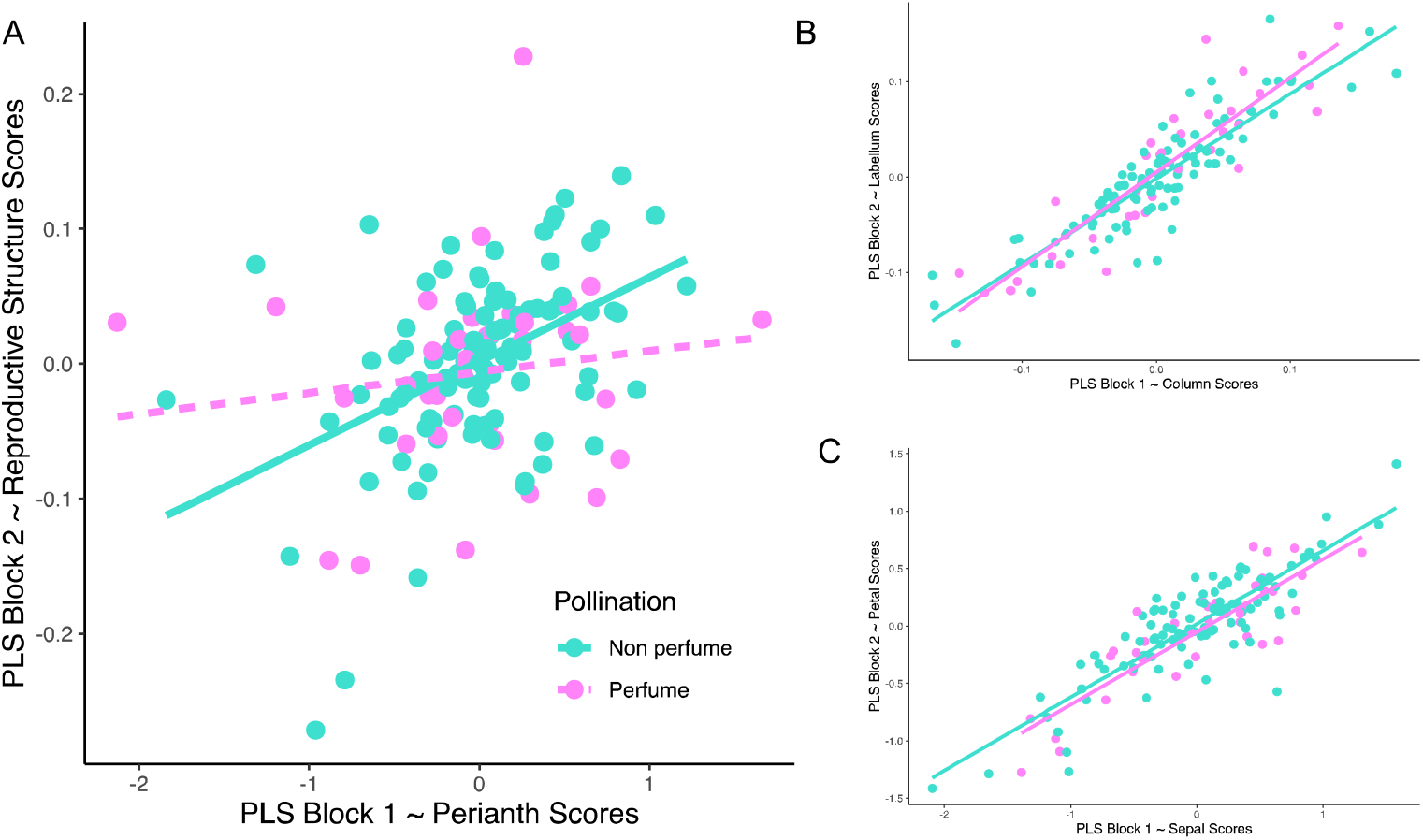
Patterns of evolutionary integration in Cymbidieae flowers. A. Comparison of integration between the perianth and reproductive structures. B. Comparison of integration within the perianth. C. Comparison of integration within the reproductive structure

The primary axis of variation in the perianth dataset explaining almost 50% of the total variance captures elongation (Fig. 1B). Species with high values of PC1 exhibit exaggeratedly long petals and sepals, as exemplified by the “spider orchid” *Brassia caudata*, while those with low values along this axis of variation exhibit squat, rounded floral organs (Table S1). The second PC, explaining around 20% of morphological variation, contrasts species with long petals relative to the sepals or vice versa. Mirroring the reproductive structure dataset, this axis roughly divides the morphospace between the diverse Maxillariinae and Oncidiinae clades (Fig. S2B).

We further categorized the pollination mechanisms of the studied species with morphological data. After excluding species with no observations or supporting evidence for pollination mode, we reduced our dataset to 140 species for further comparative analyses. Of these, 104 were classified as non-perfume flowers, spanning a continuous gradient between deceptive and rewarding taxa, and 36 were classified as perfume flowers, offering only perfumes as both signaling agents and rewards to male euglossine bees. We used stochastic character mapping to characterize the history of pollination mode evolution across the Cymbidieae phylogeny, recovering a mean of 5.16 transitions from non-perfume flowers to perfume flowers and 2.29 reversions, with lineages spending approximately three times as much evolutionary time spent in the non-perfume flower state (Table 1).

**Table 1:** Comparison of results from stochastic character mapping, morphological disparity analyses, and evolutionary rate modelling among pollination modes.

| Pollination mode (n) | Time in tree (%) | Morphological disparity | Multivariate Rate |
| --- | --- | --- | --- |
| Perfume (36) | 25.3 | 0.213 (reproductive structure)<br>11.710 (perianth) | 1.302 (reproductive structure)<br>1.633 (perianth) |
| Non perfume (104) | 74.7 | 0.173 (reproductive structure)<br>8.860 (perianth) | 0.698 (reproductive structure)<br>0.367 (perianth) |

Perfume flowers occupy an equivalent amount of morphospace in both datasets compared to flowers of all other pollination mechanisms, suggesting a strong influence on pollination mode in elevating the rates of morphological evolution (Table 1). Indeed, pollination mode had a strong effect on the rate of floral evolution, with posterior probabilities of state dependence at 0.94 and 1.0 for the reproductive structure and perianth, respectively. Critically, perfume flowers evolved at 1.9-fold and 5.3-fold the pace of non-perfume flowers for these two sets of traits (Fig. 1), accounting for their equivalent amounts of floral morphospace filled despite having a third of evolutionary time to accumulate this variation.

We detected strong integration within both structures (r-PLS > 0.8, P < 0.0001), indicating covariation between petals and sepals and between the column and labellum. We detect substantial integration between the reproductive structure and the perianth in non-perfume flowers (Z = 2.715, P = 0.005) in contrast to perfume flowers that exhibit weak integration (Z = 0.825, P = 0.216). The significance of integration in non-perfume flowers was conserved after randomly sampling them 1000 times to match the smaller sample size of the perfume flowers (median r-PLS = 0.556, Z = 1.82, P = 0.034).

## Discussion

We identified a strong effect of androeuglossophily on increasing the rate of floral shape evolution in neotropical Cymbidieae orchids. These patterns emerge against a complex background of rapid morphological evolution characteristic of one of the most spectacular radiations of flowering plants in the world’s richest botanical hotspots (Fig. S3). These elevated rates of evolution in two separate groups of floral traits align with a breakdown of their evolutionary integration, potentially due to functional decoupling in some of the most extreme pollination strategies of perfume flowers. Our results run counter to the prevailing notion that stabilizing selection, via pollinator-mediated selection, results in convergence of similar and stable suites of floral traits (3, 5, 30–32).

Due to their range of grotesque appearances and complex pollination mechanisms, perfume flowers have long captured the attention of evolutionary biologists, botanists, and horticulturists alike (23). Many species manipulate the movement of their male euglossine bee pollinators – from causing them to fall through open dimensions of the flowers (e.g. *Gongora* and *Stanhopea*) to forcing them to crawl through narrow passages to escape drowning (e.g. *Coryanthes*). Additionally, many species employ complex strategies of pollinarium placement, such as the forceful propulsion of pollen structures in *Catasetum* and the hinged labellum of *Peristeria* and *Gongora* subgenus *Acropera* causing visiting bees to be propelled into the column (24, 33). These mechanisms contrast with more passive pollination strategies found in other modes among orchids. For example, “gullet flowers” are a common feature of many members of the hyperdiverse Maxillariinae clade, which simply require crawling in and out of a passage created by the petals and sepals to ensure proper placement of pollinaria (22).

We detected 2- and 5-fold increases in evolutionary rate of the reproductive and perianth structures, respectively, in perfume flowers compared to flowers utilizing other pollination mechanisms. Notably, we found that the fastest rates of evolution in these structures occur in the Catasetinae and the Stanhopeinae, respectively, two groups of orchids that have been extensively studied in the context of male euglossine pollination. This increased pace of evolutionary change has allowed perfume flowers to accumulate equivalent amounts of morphological variation in only around a quarter of the total evolutionary time in the clade, resulting in some of the most extreme floral morphologies and pollination mechanisms in the orchids and across flowering plants.

An apparent inverse relationship between morphological and species diversity is present across the Cymbidieae. This pattern is consistent with the history of systematic revisions in the two most diverse clades of the Cymbidieae – the Maxillariinae and the Oncidiinae – that are predominately composed of non-perfume flowers (34–36). In these clades, multiple major upheavals of taxonomic organization were carried out with the advent of genetic data, showing pervasive morphological convergence in floral form, likely in response to utilization of the same guilds of pollinators within both clades (Meliponini bees and Centridini bees, respectively (37)). The presence of rapid evolution in lineages that utilize alternative strategies within each clade (e.g. *Cryptocentrum* in the Maxillariinae that is putatively moth-pollinated exhibits some of the fastest rates of labellar evolution) mirrors the patterns of release from constraint found in previous studies of pollinator-mediated morphological stasis (3).

We observed strong integration across our dataset within the perianth and reproductive structures, potentially due to shared development, coupling due to similarities in function, or a combination of both. Furthermore, we found that non-perfume flowers exhibit significant integration between their perianth and reproductive structures. We note that in the two most diverse clades containing non-perfume flowers, the Maxillariinae and the Oncidiinae, coupled function among these sets of traits is expected. For example, many species in the former group exhibit gullet flowers, forcing pollinators to contact the column through a passage created by the petals and sepals as they enter to collect rewards. In the latter group, flowers of many species mimic Malpighiaceous flowers that offer oil to female *Centris* bees, with functional coupling across the flower expected to create this visual similarity (37, 38).

By contrast, in perfume flowers we detect a lack of significant evolutionary integration between the perianth and reproductive structures, suggesting decoupling of these regions within their flowers. The role of evolutionary integration as a force promoting or hindering diversification in complex systems is controversial (27, 39, 40). However, several studies have suggested the latter in floral systems, where reduced integration (i.e., increased modularity) facilitates more rapid morphological evolution due to release from constraints. For example, increased modularity was found to increase evolutionary rates in the neotropical Melastomataceae, allowing for rapid transitions between divergent pollination modes (31). Additionally, modularity in concordance with developmental hypotheses was identified in Malagasy *Bulbophyllum* orchids, allowing for independent evolution of different floral components (32). Our data are consistent with this framework, with weak integration in the perfume flowers perhaps contributing to their exceptional diversification in floral form into extreme areas of shape space facilitating evolution of their complex pollination mechanisms.

While the arrival of Cymbidieae orchids to the Americas was coincident with the origin of perfume collection in euglossine bees (both around 34 million years ago), predominately androeuglossophilous lineages exhibit much younger ages (∼15 million years for the Stanhopeinae and the Catasetinae, with many species originating in the past 5 million years (10, 41)). Furthermore, rates of morphological evolution exhibit an acceleration towards the present, potentially facilitated by the rapid accumulation of euglossine bee species over the last 10 million years (42). The high levels of diversity in sympatric bee populations, with up to 66 species in northwestern Amazonian regions (21), may have facilitated rapid morphological evolution to minimize potential costly hybridization between interfertile orchid taxa (43).

Our results provide a new perspective on the influences of pollination on the dynamics of floral evolution. Using a series of rigorous phylogenetic comparative methods, we reconstructed the evolutionary history of pollination in the neotropical Cymbidieae radiation and characterize the substantial impact of male euglossine pollination in triggering the exceptional diversification of shape in perfume flowers. This shift in the adaptive landscape has allowed this group of plants to evolve some of the most complex pollination mechanisms that have captivated evolutionary scientists for over a century.

## Methods

### Data Collection

We collected most of our measurements from “Lankester composite digital plates” – composite arrangements of photographs from dissected flowers (Fig. S1). Additionally, we took photographs of dissected flowers taken from private and public collections as well as comparable photographs from published studies (Table S4). We conducted two sets of morphological analyses – a shape analysis using geometric morphometrics from lateral views of the column and labellum (the “reproductive structure”), and linear and curved measurements of the sepals and remaining unmodified petals (the “perianth”).

Data for the reproductive structure were collected using StereoMorph v1.6.5 (44). We placed 4 homologous landmarks corresponding to the intersection of the column with the anther cap, the intersection of the column with the ovary, the intersection of the ovary with the labellum, and the distal tip of the labellum. Three sets of curves were drawn between these points that corresponded to the column, column foot, and labellum, with 15, 25, and 10 equidistant points sampled from these regions, respectively.

For the perianth, we took 15 linear measurements per flower corresponding to 5 measurements of the organ lengths, 5 measurements of organs at their points of attachment, and 5 measurements of maximum widths across organs. Due to the presence of missing data from occasional broken and missing organs, we then averaged paired measurements (e.g. left and right petal lengths) for a total of 9 final measurements that were present in every species. These final measurements consisted of the dorsal sepal length, dorsal sepal base, dorsal sepal maximum width, lateral petal length, lateral petal base, lateral petal maximum width, lateral sepal length, lateral sepal base, and lateral sepal maximum width (Fig. S1).

Due to the limited number of replicates within species (median of 2 individuals), we used a Procrustes ANOVA to quantify the variation among individual flowers in 14 species where 4 or more flowers were photographed. We found that species contributed 94.77% of the variance, while replicate contributed only 1.66%, suggesting that our sampling methodology was appropriate for the broad scales of morphological variation we were interested in.

### Controlling for Size and Log-Shape Ratios

As the primary goal of this study was to understand shape evolution across Cymbidieae orchids, we utilized methods to remove the effect of size. For the shape analysis of the labellum and column, we performed a generalized Procrustes analysis to center and align coordinates of each specimen in a standardized shape space independent of size. As size and shape can still vary evolutionarily due to allometry, we used the *procD*.*pgls* function in geomorph v.4.0.6 (45) to perform a Procrustes ANOVA of shape vs. log(centroid-size). While the effect was significant (P = 0.023), the overall relationship was weak (r^2^ = 0.017).

We used log-shape ratios to control for size across our dataset of linear measurements. These ratios were computed by dividing each trait measurement by the geometric mean of the petal and sepal lengths that captured the major dimensions of floral size, and then taking the logarithm of this value. We then applied the *phylolm* function from phylolm v.2.6.2 to quantify the relationship between each transformed trait and log(size). The largest r^2^ value was 0.062 with no significant trends, confirming an overall weak relationship between size and shape in the transformed variables (Table S2)

### Classification of pollination modes

The neotropical Cymbidieae exhibit a diversity of pollination rewards, including the presentation of oils, nectar, resin, edible trichomes, and perfume; in addition to mimicry of those rewards and of mates (11, 29, 38). In most orchids, traits such as scent and color either signal rewards or act as cues for unrewarding flowers exploiting pre-existing mutualisms, thus mediating pollinator behavior. The distinction between these strategies is unclear – even some species that present rewards are thought to provide them in such low quantities that their validity as true mutualisms is ambiguous (35, 46, 47). In perfume flowers, however, scent acts as both the signal and reward and is thus functionally distinct from all other pollination systems.

We utilized a large published dataset of orchid pollination and rewards (29) to code pollination modes. As the majority of species did not have direct pollinator observations, we used several proxies to code pollination state. First, any species that contained nectar, trichome, oil, or resin rewards were characterized as non-perfume flowers, as by definition perfume flowers do not produce these additional rewards. Second, if at least two members of a genus produced the same reward or had multiple observations of the same pollination type, we considered congeners without those standards of evidence to exhibit the same pollination type. As “rewardless” orchids could plausibly represent either perfume flowers or deceptive pollination, we elected to not code orchids as either system if they simply were labelled as not producing rewards and did not have direct observations. After removing species with no observations or evidence of rewards, we reduced our dataset to 140 species for further comparative analyses.

### Phylogenetic comparative methods

We used the time-calibrated phylogenetic hypothesis proposed in (10) pruned to our focal species. Where sample species in our dataset were not present in the phylogeny, we substituted them with a congener or if possible, a member of the same subgenus (Table S3). To understand the evolutionary history of pollination mode evolution in the Cymbidieae, we used the *make*.*simmaps* function in phytools version 2.0 to generate 1000 stochastic character maps (48, 49). As the neotropical Cymbidieae orchids originated from a dispersal event from a continent where euglossine bees are not present (10), we fixed the root of the tree to have the non-perfume flower character state.

We used phylogenetic principal components analysis (pPCA) to visualize and understand major patterns in our trait spaces (50). For the reproductive structure dataset, we performed pPCA on the Procrustes transformed coordinates using the *gm*.*prcomp* function in geomorph v.4.0.6. For the perianth dataset, pPCA was performed using the correlation rather than covariance matrix among the variables, ensuring more equal contribution of each of the variables that originally exhibited contrasting levels of variance. Though there were nine total principal component scores, only the first eight scores portrayed floral variation, since one degree of freedom was lost when applying log shape-ratios. To quantify occupation of morphological space by orchids of each pollination mode, we used the *morphol*.*disparity* function in geomorph v.4.0.6 to compare their levels of shape variance.

We utilized a Multiple State-Specific Rates of continuous-character evolution model (51) to understand the effect of pollination on floral morphological diversification. In contrast to other methods developed for continuous trait evolution, this Bayesian approach explicitly models background rate variation, a feature we expect to be present across this hyperdiverse radiation, to robustly estimate rate parameters associated with discrete states. Furthermore, this framework more accurately captures the multivariate nature of our morphological datasets. The Monte Carlo Markov Chain (MCMC) ran for 500,000 generations for the first 5 PC’s of labellum-column shape explaining 90% of variation and for 1,000,000 generations for the log-shape ratio dataset, with a prior of 4 rate shifts specified. Due to strong covariance between lateral petal and lateral sepal lengths in the shape ratio dataset leading to issues with estimations of correlation structure and model convergence, we removed the latter trait. To assess the effects of specifying different number of rate shifts, we performed both sets of analyses using priors of 1, 4, and 10 rate shifts and identified similar qualitative results (Fig. S4). We ran the model in Revbayes version 1.2.1 (52) and used Tracer version 1.7.2 (53) to verify convergence of the MCMC.

### Integration

We used phylogenetic partial least squares (pPLS) to identify patterns of covariation within and among organs in our dataset (54). For the perianth dataset, we used standardized measurements of the log-shape ratios described above while Procrustes transformed coordinates were used for the reproductive structure dataset. We used three sets of integration tests – 1) within the perianth, computing integration between the petal and sepal measurements, 2) within the reproductive structure, computing integration between the column and labellum coordinates, and 3) between the perianth and reproductive structure. To understand whether pollination mode was associated with differences in integration, we computed these values within the perfume flowers and non-perfume flowers. All integration analyses were performed using the *phylo*.*integration* function in geomorph v.4.0.6.

As there were far fewer perfume flowers (36) than non-perfume flowers (114) in our dataset, we randomly rarefied the non-perfume flower dataset to the size of the perfume flower dataset 1000 times. We then performed the analyses above on the 1000 balanced datasets to assess whether differences in detected integration could solely be attributed to differences in sample size.

## Acknowledgements

The authors are grateful to Guido Deburghgraeve, Danilo Zavatin, Gustavo Montealegre, and Fabio Romero for providing additional photographs of several species of orchids. We are grateful to Andy Phillips, Marni Turkel, Jeff Tyler, and the UC Davis Conservatory for allowing us access to their collections for sampling of flowers. We are thankful for the helpful feedback provided by the Ramírez Lab at earlier stages of the study and to Peter C. Wainwright and Collin P. Gross for helpful comments on the initial draft of the manuscript.

