## Supplementary information for "Exceptional diversification of floral form in a specialized orchid pollination system"

Table S1: Loadings of each principal component of perianth variation. Bolded values correspond to loadings of greater magnitude than 0.3

| | **PC1** | **PC2** | **PC3** | **PC4** |
|---|---|---|---|---|
| Dorsal sepal length | 0.128 | **0.495** | **0.851** | 0.0496 |
| Dorsal sepal base | **-0.815** | 0.162 | 0.171 | **-0.325** |
| Dorsal sepal max width | **-0.894** | 0.0132 | 0.0252 | **0.313** |
| Lateral petal length | -0.221 | **0.969** | -0.0318 | -0.0638 |
| Lateral petal base | **-0.759** | 0.0558 | -0.102 | **-0.512** |
| Lateral petal max width | **-0.882** | -0.0868 | -0.0654 | 0.225 |
| Lateral sepal length | 0.133 | **0.644** | **-0.751** | 0.0267 |
| Lateral sepal base | **-0.789** | 0.243 | 0.0243 | -0.122 |
| Lateral sepal max width | **-0.893** | 0.0683 | -0.0318 | **0.331** |

Table S2: Results of phylogenetic regression of log(size) on log shape ratios for each perianth trait

| Trait | Coefficient | P-value | $R^2$ |
|---|---|---|---|
| Dorsal sepal | 0.02200 | 0.28960 | 0.00246 |
| Dorsal sepal base | -0.12836 | 0.30491 | 0.06059 |
| Dorsal sepal width | -0.03564 | 0.72838 | 0.02645 |
| Average petal base | -0.10333 | 0.32360 | 0.04285 |
| Average lateral sepal base | -0.12982 | 0.29216 | 0.06063 |
| Average petal width | -0.10457 | 0.35188 | 0.02448 |
| Average lateral sepal width | -0.01092 | 0.91981 | 0.02346 |
| Average petal length | -0.01488 | 0.49970 | 0.01003 |
| Average lateral sepal length | -0.00712 | 0.71350 | 0.00461 |

Table S3: Substitutions of species made in this study

| Sampled species | Substituted species in phylogeny |
|---|---|
| *Brasiliorchis picta* | *Brasiliorchis picta* |
| *Brassia ocanensis* | *Brassia allenii* |
| *Camaridium bracteatum* | *Camaridium bradeorum* |
| *Catasetum planiceps* | *Catasetum collare* |
| *Catasetum maculatum* | *Catasetum expansum* |
| *Catasetum saccatum* | *Catasetum macrocarpum* |
| *Chondrorhyncha chocoensis* | *Chondrorhyncha rosea* |
| *Chondroscaphe chestertonii* | *Chondroscaphe amabilis* |
| *Christensonella uncata* | *Christensonella echinophyta* |
| *Cischweinfia parva* | *Cischweinfia colombiana* |
| *Cochleanthes aromatica* | *Cochleanthes flabelliformis* |
| *Coryanthes hunteriana* | *Coryanthes elegantium* |
| *Cryptocentrum latifolium* | *Cryptocentrum calcaratum* |
| *Cycnoches warcswiczii* | *Cycnoches cooperi* |
| *Cyrtidiorchis gerardii* | *Cyrtidiorchis alata* |
| *Cyrtopodium macrobulbon* | *Cyrtopodium aliciae* |
| *Dichaea andina* | *Dichaea pendula* |
| *Galeandra arundinis* | *Galeandra minax* |
| *Gongora claviodora* | *Gongora ilense* |
| *Heterotaxis crassifolia* | *Heterotaxis sessilis* |

| | |
|---|---|
| *Houlletia lowiana* | *Houlletia sanderi* |
| *Ixyophora velatiguii* | *Ixyophora viridisepala* |
| *Kefersteinia alata* | *Kefersteinia excentrica* |
| *Lacaena spectabilis* | *Lacaena bicolor* |
| *Macroclinium lineare* | *Macroclinium aurorae* |
| *Miltonia moreliana* | *Miltonia candida* |
| *Mormodes lobulate* | *Mormodes andreettae* |
| *Oliveriana uribe-velezii* | *Oliveriana brevilabia* |
| *Paphinia cristata* | *Paphinia neudeckeri* |
| *Pityphyllum mercedes-abarcae* | *Pityphyllum huancabambae* |
| *Psychopsis krameriana* | *Psychopsis limminghei* |
| *Rodriguezia pulcherrima* | *Rodriguezia batemanii* |
| *Sauvetrea sessilis* | *Sauvetrea chicana* |
| *Scuticaria steelei* | *Scuticaria hadwenii* |
| *Stanhopea wardii* | *Stanhopea saccate* |
| *Vargesiella colombiana* | *Vargasiella peruviana* |
| *Zygopetalum crinitum* | *Zygopetalum maculatum* |

Table S4: Sources of additional species not sampled in this study

| **Species** | **Source** |
|---|---|
| *Acineta superba* | Sierra-Ariza et al. 2022 (1) |
| *Clowesia dodsoniana* | Tamayo-Cen et al. 2022 (2) |
| *Clowesia russelliana* | Tamayo-Cen et al. 2022 (2) |
| *Cycnoches egertonianum* | Romero-González 2018 (3) |
| *Cycnoches warscewiczii* | Romero-González 2018 (3) |
| *Cyrtidiorchis gerardii* | Uribe-Velez et al. 2020 (4) |
| *Cyrtochilum myanthum* | Morales et al. 2018 (5) |
| *Cyrtochilum ramosissimum* | Dalstrom 2020 (6) |
| *Dichaea hystricina* | Moreno et al. 2020 (7) |
| *Gomesa longipes* | Davies et al. 2014 (8) |
| *Gomesa radicans* | Stpiczyńska and Davies, 2008 (9) |
| *Houlletia lowiana* | Sierra-Ariza et al. 2022 (1) |
| *Huntleya wallisii* | Uribe-Velez and Sauleda, 2020 (10) |
| *Ixyophora velatiguii* | Pupulin 2019 (11) |

| | |
|---|---|
| *Oliveriana uribe-velezii* | Sauleda and Uribe-Velez, 2020 (12) |
| *Oncidium alexandrae* | Karremans et al. 2017 (13) |
| *Oncidium fuscatum* | Sierra-Ariza et al. 2022 (1) |
| *Oncidium portilloides* | Uribe-Velez and Sauleda, 2020 (14) |
| *Ornithocephalus escobarianum* | Moreno et al. 2020 (7) |
| *Pityphyllum mercedes-abarcae* | Vélez et al. 2021 (15) |
| *Sauvetrea sessilis* | Sierra-Ariza et al. 2022 (1) |
| *Trichocentrum jonesianum* | Carnevali et al. 2010 (16) |
| *Vargasiella colombiana* | Karremans et al. 2017 (13) |
| *Vitekorchis excavata* | Davies et al. 2014 (8) |

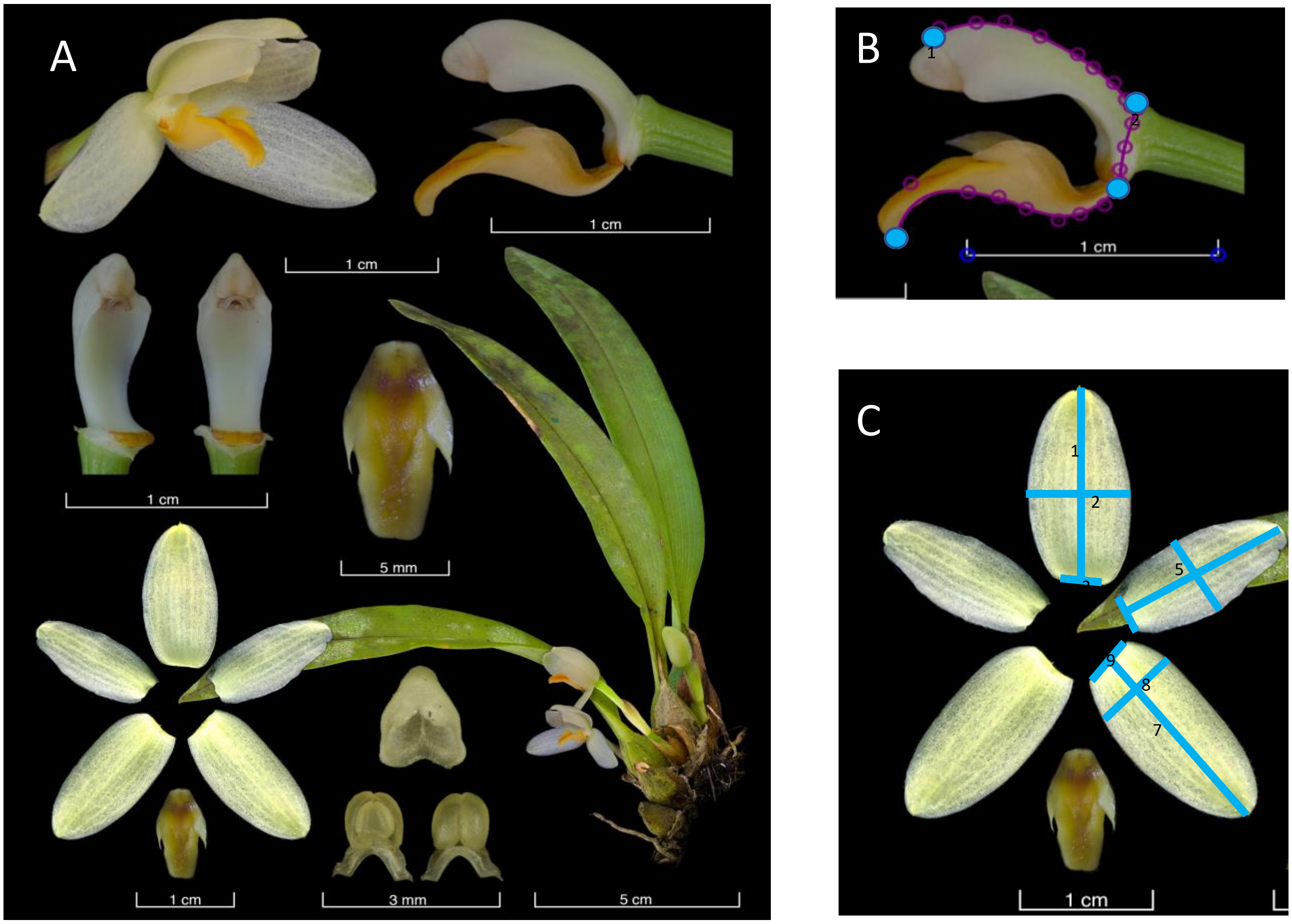


Figure S1: Example of images used in this study. A. A Lankester composite digital plate of *Mormolyca hedwigiae*. B. Landmarking scheme of the reproductive structure. Landmarks consist of 1) Anther cap column junction, 2) column-ovary junction, 3) column foot-labellum junction, and 4) distal tip of the labellum C. Linear measurements taken of the perianth. Labelled measurements correspond to 1) length of the dorsal sepal, 2) maximum width of the dorsal sepal, 3) width at the base of the dorsal sepal, 4) length of the lateral petal, 5) maximum width of the lateral petal, 6) width at the base of the lateral petal, 7) length of the lateral sepal, 8) maximum width of the lateral sepal, and 9) width at the base of the lateral sepal

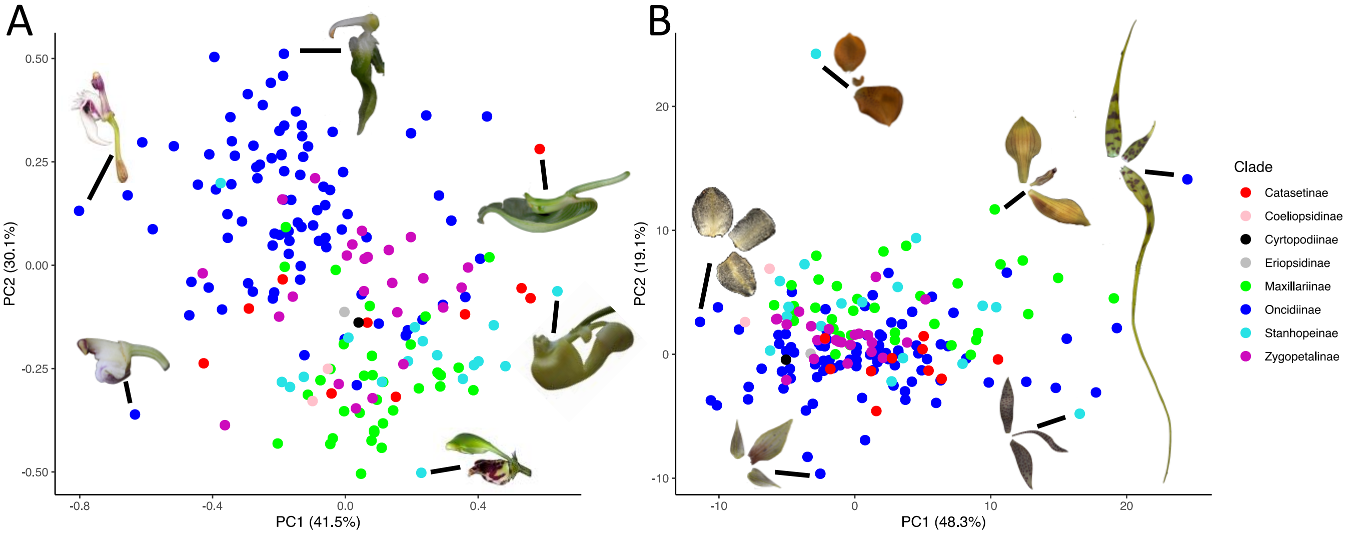


Figure S2: Morphospaces of the neotropical Cymbidieae flower comparing axes of variation in A. the reproductive structure and B. the perianth. Points represent species means and are colored by clade. Other represented species in A are (from center above, going counterclockwise): *Ornithocephalus inflexus, Cyrtochilum ramosissimum*, *Cyrtochilum myanthum*, *Kegeliella kupperi*, *Coryanthes hunteriana*, and *Clowesia dodsoniana*. Other represented species in B are (from center above, going counterclockwise): *Gongora armeniaca*, *Trichocentrum morenoi*, *Telipogon biolleyi*, *Polycycnis barbata*, *Brassia caudata*, and *Trigonidium egertonianum*. Photographs copyright the authors except for the following, modified with permission: *Clowesia dodsoniana* from (2), *Cyrtochilum ramosissimum* from (6) copyright Carlos Jerez, *Cyrtochilum myanthum* from (5), and *Trichocentrum morenoi* copyright Gustavo Montealegre.

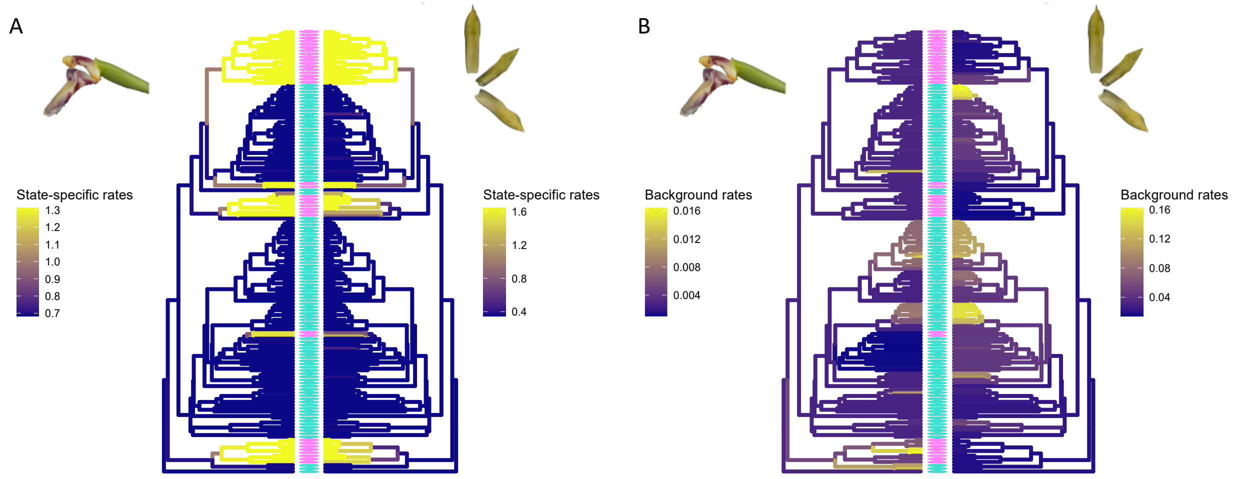


Figure S3: State-specific (A) and background rates (B) of morphological evolution across the Cymbidieae. Branch colors on the phylogenies indicate respective rates of evolution, with dark purple indicating lower rates and yellow indicating higher rates. Bars between the colored phylogenies represent character states at the tips, with pink corresponding to perfume flowers and turquoise corresponding to non perfume flowers.

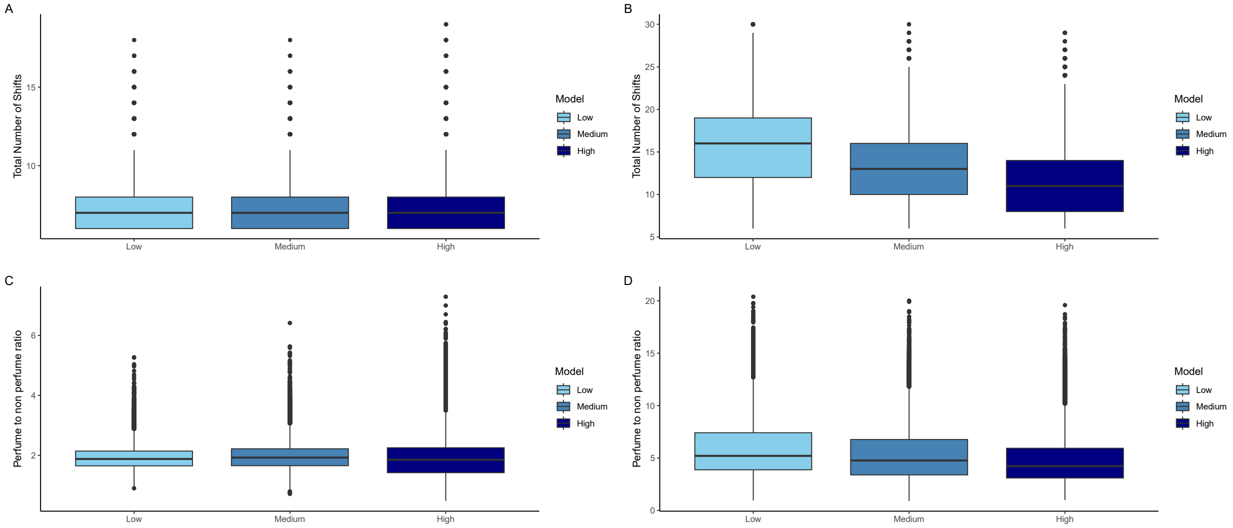


Figure S4: The influence of different rate shift priors (1, 5, and 10 for low, medium, and high, respectively) on total number of shifts (A and B) and the ratio of rates of evolution of perfume flower to non perfume flowers (C and D). While the number of rate shifts decreases with increased priors, the ratio of rates of morphological evolution between perfume and non perfume remains similar.
